# Digital Immunoassay for Rapid Detection of SARS-CoV-2 Infection in a Broad Spectrum of Animals

**DOI:** 10.1101/2024.06.21.600129

**Authors:** Siyan Li, Weijing Wang, Weinan Liu, Chi Chen, Skye Shephard, Fangfeng Yuan, Jennifer M. Reinhart, Diego Diel, Brian T. Cunningham, Ying Fang

**Affiliations:** Department of Pathobiology, University of Illinois at Urbana-Champaign, Urbana, IL, USA; Department of Bioengineering, University of Illinois at Urbana-Champaign, Urbana, IL, USA; Department of Electrical and Computer Engineering, University of Illinois at Urbana-Champaign, Urbana, IL, USA; Department of Veterinary Clinical Medicine, University of Illinois at Urbana-Champaign, Urbana, IL, USA; Department of Population Medicine and Diagnostic Sciences, Cornell University, Ithaca, New York, USA; Carl R. Woese Institute for Genomic Biology, University of Illinois at Urbana-Champaign, Urbana, IL, USA

**Keywords:** SARS-CoV-2 detection, photonic crystal biosensor, monoclonal antibody, blocking biosensor assay

## Abstract

The ability of Severe Acute Respiratory Syndrome Coronavirus 2 (SARS-CoV-2) to infect a wide-range of species raises significant concerns regarding both human-to-animal and animal-to-human transmission. There is an increasing demand for highly sensitive, rapid, and simple diagnostic assays that can detect viral infection across various species. In this study, we developed a biosensor assay that adapted a monoclonal-antibody (mAb)-based blocking ELISA format into an Activate Capture + Digital Counting (AC + DC)-based immunoassay. The assay employs a photonic crystal (PC) biosensor, gold-nanoparticle (AuNP) tags, SARS-CoV-2 nucleocapsid (N) protein, and specific anti-N mAb to detect antibody responses in animals exposed with SARS-CoV-2. We demonstrated a simple 2-step 15-min test that was capable of detecting as low as 12.5 ng of antibody in controlled standard serum samples. Based on an evaluation of 176 cat serum samples with known antibody status, an optimal percentage of inhibition (PI) cut-off value of 0.588 resulted in a diagnostic sensitivity of 98.3% and a diagnostic specificity of 96.5%. The test is highly repeatable with low variation coefficients of 2.04%, 2.73%, and 4.87% across different runs, within a single run, and on a single chip, respectively. The test was further employed to detect antibody responses in multiple animal species as well as investigate dynamics of antibody response in experimentally infected cats. This test platform provides an important tool for rapid field surveillance of SARS-CoV-2 infection across multiple species.

## INTRODUCTION

The global outbreak of Coronavirus Disease 2019 (COVID-19) has been primarily driven by the rapid and widespread transmission of SARS-CoV-2^1^. This virus has ability to infect an extensive array of species, leading to significant concerns regarding cross-species transmission, including both from humans-to-animals and animal-to-human^23^. The situation is further complicated by the rapid mutations of viral RNA genome, which occur as it crosses between different hosts, potentially leading to new, more virulent strains that could undermine the current control and prevention measures^4^.

One of the key strategies for disease control and prevention is the early and rapid identification of infected individuals and animals, which requires highly sensitive and rapid diagnostic assays. For SARS-CoV-2 that is able to infect multiple species, the assays must be capable of detecting the virus across a variety of species in order to timely isolation of affected individuals/animals to prevent further spreading of the virus. Laboratory diagnostic tests for SARS-CoV-2 fall into four main categories: virus isolation, viral antigen detection, molecular (viral nucleic acid) diagnostics, and host antibody detection, each serving a unique purpose in identifying the virus infection. Serological assays, especially enzyme linked immunosorbent assay (ELISA), are commonly utilized to detect specific host antibodies against SARS-CoV-2, providing insights into past virus exposure^5^. While indirect ELISA is highly sensitive and specific, its dependency on species-specific secondary antibodies limits its ability to detect antibodies across a diverse range of hosts^6^. The monoclonal antibody (mAb)-based blocking ELISA (bELISA) stands out for its increased specificity in detecting antibody responses from a broad spectrum of host species ^6–8^. The bELISA incorporates mAb reagent to recognize the specific conserved epitope in a viral antigen. The test readout is based on the mAb binding ability; thus, species-specific secondary antibody is not required. Such a test platform will allow the assay applicable to all animal species and humans.

A drawback of ELISAs is their requirement of laboratory settings, including large equipment, complex workflows, and lengthy sample-to-answer time. Recently, gold nanoparticle (AuNP)-based assays have emerged as a promising alternative to ELISA, offering a more rapid, highly sensitive and cost-effective solution^7,9,10^. Here, we utilize a novel form of biosensor microscopy in which a Photonic Crystal (PC) surface is utilized in combination with a portable and inexpensive detection instrument for quantitative detection of gold nanoparticle (AuNP) tags. In our “blocking biosensor assay” each AuNP represents one antigen-antibody binding interaction that can be digitally counted. The assay is an alternative to ELISA and b-ELISA that does not require enzymatic amplification, thus providing an immediate readout without the need for enzyme-substrate interactions to generate colored or fluorescent products.

A PC is a sub-wavelength grating structure consisting of a periodic arrangement of a low refractive index material coated with a high refractive index layer. At a particular resonant wavelength complete interference occurs, and no light is transmitted, resulting in nearly 100% reflection efficiency. The resonant reflectance magnitude is modulated by the addition of a light-absorbing AuNP upon the PC surface, resulting in a large reduction in the reflected intensity. By measuring the reflected intensity from a red LED on a pixel-by-pixel basis across the PC, images of attached nanoparticles may be gathered by illuminating the structure with collimated light through the transparent substrate, while the top surface of the PC is immersed in aqueous media. The biosensor is fabricated by a nano-replica molding approach on glass wafers and plastic films as described by our earlier publications^11–15^ . The device has a linear grating surface structure (period = 400 nm, depth = 120 nm), producing a resonant narrowband optical reflection at λ=625 nm. The structures are manufactured by an external vendor (Moxtek, Provo, UT) and fabricated to our performance specifications.

Recently, our group developed a technique called Photonic Resonator Absorption Microscopy (PRAM) that can visualize individual AuNP tags on a PC surface through resonance coupling^6^. As described in prior publications^9,10,16–20^, the detection principle of PRAM utilizes the resonant PC reflection at a wavelength of λ = 625 nm to provide a high reflected intensity from collimated low intensity LED illumination of the same wavelength into a webcam-variety image sensor. The AuNPs are strategically selected to provide strong absorption by localized surface plasmon resonance at the same wavelength^21–23^. Thus, each surface-bound AuNP registers in the PC reflected image as a location with reduced intensity, compared to the surrounding regions without AuNPs. By immobilizing target-activated AuNP probes on a PC surface, PRAM has been used to quantify nucleic acids and proteins with single-particle _resolution_18,19,24–26.

PRAM imaging offers an attractive alternative to detection of fluorophore tags, as PRAM requires only low power broadband illumination with an LED, does not suffer from photobleaching, and can provide time-course data. Unlike fluorescence-based sensing, PRAM requires only a webcam-quality image sensor that can measure reduction in reflected intensity caused by surface-attached AuNPs, and can gather images from a large field of view. We recently designed a portable version of the PRAM instrument with a cost of components totaling $7,000^27^ and current research efforts are directed towards a miniaturized PRAM instrument with a cost of $500.

In this study, we integrated the bELISA test platform with the AuNP detection system to create a novel blocking biosensor assay, which merges the advantages of both methodologies to generate a new serological detection technology. This novel approach presents an advanced tool for the quick identification of coronavirus infections. This advancement not only enhances our diagnostic capabilities but also significantly contributes to our collective efforts in understanding and controlling the cross-species transmission of SARS-CoV-2 infection.

## MATERIALS AND METHODS

### Preparation of antigen and monoclonal antibody

The nucleocapsid (N) protein of SARS-CoV-2 was expressed in *E. coli* as a 6x His-tagged recombinant protein. The N antigen was produced and purified using the methods described in our previous study^7,9^. Purified protein was dialyzed using 1x phosphate-buffered saline (PBS) solution under 4℃ and then concentrated by polyethylene glycol 8000 (Thermo Fisher Scientific, 320 Waltham, MA). The SARS-CoV-2 N protein specific mAb #127-3 was generated in our previous study^6^.

### Synthesis of Spiky Core-shell AUNPs

The spiky core-shell Gold Nanoparticles (AuNPs) seeds were synthesized using a two-step growth method. Initially, 25 µL of a commercially obtained 20-nm nanoparticle seed solution (Cytodiagnostics , 3 M) was incubated in 600 µL of a 20 mM sodium citrate solution for 30 minutes with moderate shaking. After incubation, the nanoparticles were washed by centrifugation at 20,000 g once and then re-dispersed in 2 mL of a 6 mM sodium citrate solution (Sigma-Aldrich). Next, 4 mL of a 0.5 mM HAuCl solution (Sigma-Aldrich) was brought to a boil, and 2 mL of the nanoparticle seed solution was rapidly added while stirring vigorously at 700 rpm. The mixture was allowed to react for 10 minutes at boiling temperature, during which the solution color changed from brown to burgundy. DI water was added during the process to maintain a consistent solution volume. The solution was then cooled under stirring for another 10 minutes, followed by centrifugation at 4,000 g three times. The resulting smooth spherical core-shell AuNPs were re-dispersed in 1.5 mL of a 2 mM sodium citrate solution to serve as the seeds for the next step. For the growth of the spiky gold coating, a mixture of 5 mL of 0.25 mM HAuCl, NP seeds (Absorbance at 535 nm = 3.0, 100 µL), and 11 µL of 1% (w/v) sodium citrate was stirred vigorously. Subsequently, 500 µL of a 30 mM hydroquinone solution (Sigma-Aldrich) was quickly injected to initiate the reaction. The growth solution was allowed to react for 30 minutes at room temperature under consistent stirring, during which the solution gradually changed from pink to blue. The resulting spiky AuNP solution was washed three times at 1,000 g and re-dispersed in a 0.1% (w/v) Pluronic F-127 solution for long-term storage.

### Anti-N mAb-AuNP conjugation

Initially, a fresh aliquot of SH-PEG-NHS (MW 5000, JenKem Technology, Plano, TX, USA) is diluted in Milli-Q water to a final concentration of 4 µM. Subsequently, 20 µg of the anti-SARS-CoV-2 N mAb is gently mixed with the SH-PEG-NHS solution at room temperature, using a pipette to ensure thorough integration without vigorous agitation. This mixture is then incubated on a shaker at 900 rpm for 2 hours to facilitate the conjugation of the SH-PEG to the mAb. Following the conjugation, the mixture is filtered using a centrifuge filter (MilliporeSigma™ Amicon™ Ultra-0.3 Centrifugal Filter Units) to remove any unbound PEG. This step is performed at 4°C to preserve the integrity of the proteins and linker. The concentrated SH-PEG-mAb conjugate is then recovered and quickly combined with AuNPs in a non-stick microcentrifuge tube, avoiding any form of vortexing to prevent damage to the delicate conjugates. The AuNPs and SH-PEG-mAb are then left to incubate overnight at 4°C, allowing the mAb to bind efficiently to the AuNPs. Post-incubation, the conjugates undergo Dynamic Light Scattering (DLS) analysis to verify the increase in particle size, indicative of successful conjugation. The solution is then stabilized by adding filtered 2% BSA and further incubated to ensure complete coating of the conjugates. Excess mAbs are removed by centrifugation (1000 rcf, 5 mins), and the conjugates are resuspended in a 10% PBST (Pierce™) solution for final storage at 4°C, where they remain stable for up to one week. This preparation is critical for subsequent applications in active capture and digital counting assays, where the functionality and stability of the conjugates directly influence the assay’s sensitivity and specificity.

### PC biosensor preparation and functionalization

For the preparation of photonic crystal (PC; Moxtek, Orem, UT), the process begins with the proper assembly of the PC on a coverslip using a droplet of UV glue (Norland Optical Adhesive NOA 68, 1 oz), followed by UV light curing for a minimum of two minutes to secure the attachment. To determine the correct orientation, the PC’s edges are tested with tweezers; the side that scratches easily is identified as the top. Subsequent cleaning involves immersing the PC in a glass jar filled with acetone and utilizing a sonicator bath for two minutes. This process is repeated with isopropyl alcohol (IPA) and Milli-Q water to ensure thorough cleansing. The PC is then dried under a stream of nitrogen and placed in a 60°C oven to evaporate residual moisture. Surface activation of the PC is conducted using an oxygen plasma generator set at 100% power for 10 minutes. Concurrently, a 5% solution of 3-Aminopropyltriethoxysilane (APTES, Sigma-Aldrich) in tetrahydrofuran (THF, Sigma-Aldrich) is prepared by mixing 47.5 mL of THF with 2.5 mL of APTES. The activated PC is transferred into this APTES solution, where it is incubated for one hour on a shaker set at 600 rpm and room temperature. Following incubation, the APTES solution is discarded, and the PC is further cleaned using THF, acetone, ethanol, and Milli-Q water in a sonication process. Finally, the PC is dried, wrapped in aluminum foil to protect it from light, and stored in a desiccator for up to one month, ensuring its readiness for subsequent functionalization and assays.

### Sample sources

The control serum standards utilized for assay development were prepared using samples obtained from our previous cat experiment. The positive control standard was made from serum samples of experimental cats that were infected with SARS-CoV-2 variants D614G (B.1), Delta (B.1.617.2), and Omicron (B.1.1.529) at 14 days post-infection (dpi). Similarly, the negative control standard was made from serum samples of negative control cats. Large volumes of positive sera from different cats were pooled into a single lot of positive control standard, while large volumes of negative sera were pooled into a single lot of negative control standard.

To apply the blocking biosensor assay in the diagnosis of clinical animals, 186 serum samples were obtained from Clinical Pathology Laboratory of the University of Illinois Veterinary Diagnostic Laboratory in Urbana, IL. These samples were collected from excess (leftover) serum samples that were originally submitted for other clinical purposes. Since the serum was not collected purposely for this study, the sample collection was not qualified as animal use under the regulations, which was not required the approval from the University of Illinois Institutional Animal Care and Use Committee^28^. Furthermore, to ensure confidentiality, all identifiable information, such as animal names and addresses, was removed from the data, and the new project specific identifiers were assigned to each sample.

For validation of the biosensor assay in multiple animal species, two sets of animal serum samples with known infection status were used. The first set contained 5 positive and 3 negative serum samples collected from ferrets experimentally infected by SARS-CoV-2 isolate NYI67-20 (B.1 lineage). The second set contained 5 positive and 3 negative serum samples collected from the deer infected by SARS-CoV-2 isolate SARS-CoV-2 isolate NYI67-20 (B.1 lineage). To apply the blocking biosensor assay for detecting the seroconversion, a panel of serum samples were collected from 3 experimental cats in a time course study at 0, 3, 7, 10, and 14 days post-inoculation. Among these 3 cats, one was infected with SARS-CoV-2 Omicron (B.1.1.529) variant, and another one was infected with D614G (B.1) variant, while samples from one negative control cat were included for comparison.

### Procedure for blocking biosensor assay

Initially, the PC was initially coated with 10 μL of N protein (8 μg/mL) and left to incubate overnight at 4°C. Subsequently, each well was treated with 20 μL of 1x power block (Thermo fisher) and incubated at 37°C for 1 hour to significantly reduce the nonspecific binding of AuNPs during the assay. Following this, the PC underwent a washing process three times with 1x PBST. Finally, serum samples, diluted 4x with blocking buffer, were applied to the PC surface and incubated at 37°C for a duration of 1 hour on a shaker to ensure comprehensive binding. The AuNPs-PEG-mAb conjugate is sonicated for 10 seconds to ensure homogeneity and checked for the absence of aggregates using Dynamic Light Scattering (DLS).

For the assay setup, the PC coated is placed under the Photonic Resonator Absorption Microscopy (PRAM) instrument. A background image is captured after ensuring the PDMS well is clear and free of any obstructions. Then the sample serum in each PDMS wells will be removed by pipetting. Following this, 20 µL of AuNPs-PEG-mAb conjugates will be added to the PDMS well. The reaction mixture is then left to react for 15 minutes. Afterward, the PC is aligned under the PRAM to capture the reaction image. This process is repeated for each concentration, with each sample tested at least two times to ensure reliability. The images obtained are processed using a specific Matlab algorithm, which adjusts the threshold and other parameters to accurately count the particles, ensuring that the counting does not include background noise or irrelevant data. This detailed approach helps quantify the presence of Anti-SARS-CoV-2 N mAbs effectively.

## RESULT

### Photonic crystal biosensor design and functionalization

The PC biosensors were fabricated on 8-inch diameter glass substrates based on our designed specifications and cut into 10 × 12 mm^2^ pieces for use in all subsequent experiments. The PC has a linear grating period of 380 nm and a grating depth of 97 nm etched into a glass substrate by reactive ion etching. The grating is coated by TiO_2_ thin film with a high refractive index of 2.25 and a thickness of 98.5 nm, which yields a surface structure with high-efficiency resonant reflection at a wavelength of 625 nm when the PC surface is covered in aqueous media. PCs are used once and are discarded after an assay.

### Preparation and characterization of anti-N mAb-AuNP conjugate

We prepared Anti-N mAb-AuNPs by conjugating Anti-N mAb#127-3 with 90 nm diameter gold nanourchins via heterobifunctional SH-PEG-NHS linkers (Fig. 1A). Gold nanourchins with a protruding tip morphology were chosen as the label for PRAM-based imaging due to their enhanced light harvesting across the particle surface, and their surface plasmon resonance wavelength also closely matches the PC resonant reflection wavelength. The prepared mAb-AuNPs were characterized by UV-Vis spectroscopy to confirm the conjugation of the Anti-N mAb. A slightly red-shift (∼5 nm) was observed after the mAb modification and the maximum absorption wavelength is approximately 625 nm, which perfectly matches the resonant reflection wavelength of the PC. As a gauge of conjugation performance, dynamic lighting scattering (DLS) measurements showed that mAb-AuNPs displayed a distinct increase of ∼ 25 nm in average diameter compared to non-functionalized bare AuNPs (110 nm vs 136 nm).

**Figure 1.**
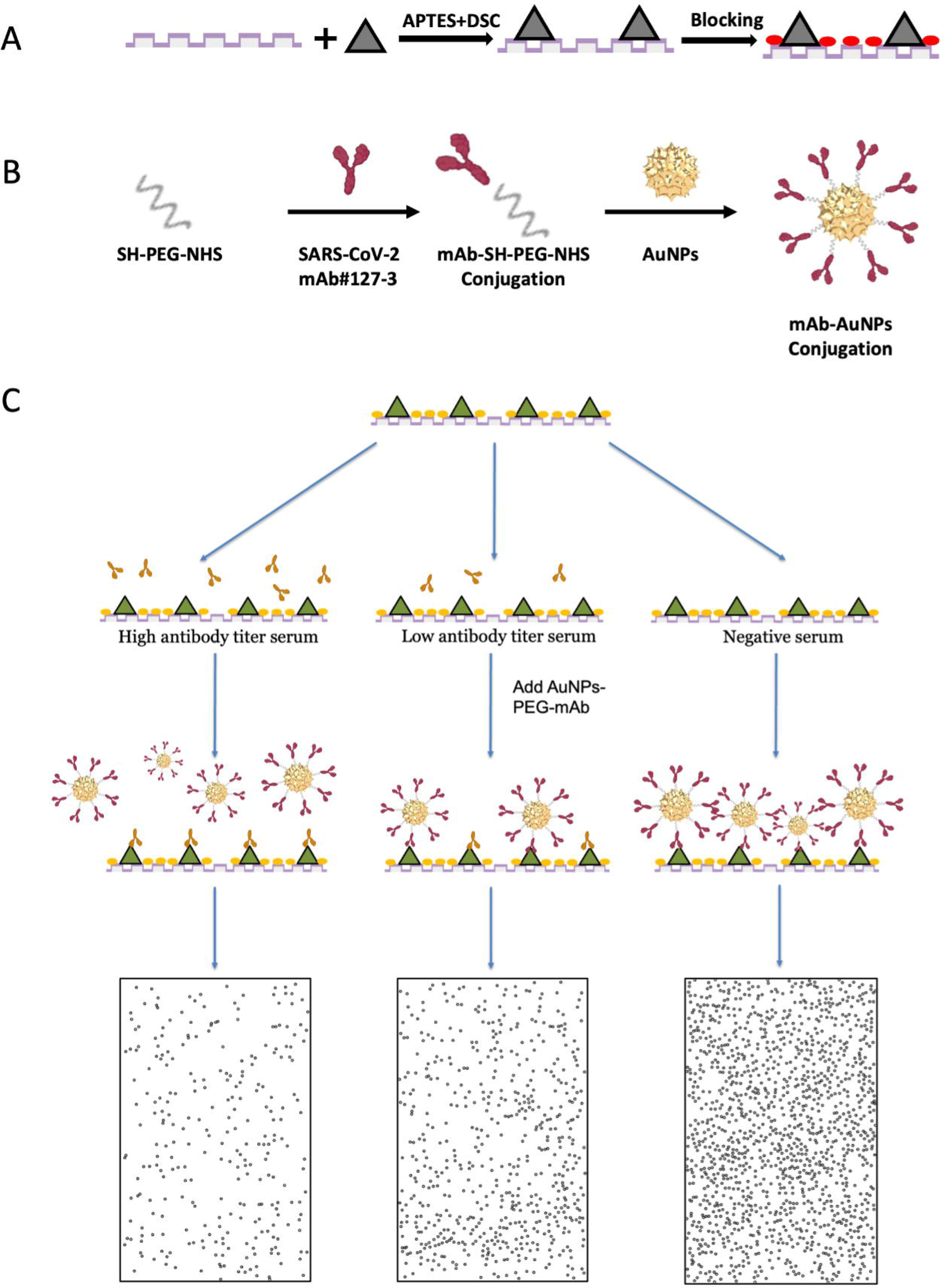
Schematic illustration of Blocking Biosensor immunoassay for detection of host antibodies against SARS-CoV-2. **(A) (B)** Preparation of monoclonal antibody functionalized AuNPs. Anti-N monoclonal antibodies were conjugated with 80 nm diameter gold nanourchins via SH-PEG-NSH linkers. (**C**) Serum sample was added to a PC surface coated with N protein, allowing anti-N antibodies in the serum to bind. AuNPs-conjugated mAbs were then added to react with any unbound N protein. PRAM image peak intensity values varied with the concentration of AuNPs-mAb on the PC surface. Higher anti-N antibody titers in the serum reduced mAb-AuNPs binding, decreasing the AuNP count observed under PRAM microscopy. The assay took 15 minutes.

### Design of mAb-based blocking biosensor digital immunoassay

This assay adopts the same immunological structure as the bELISA but incorporates PC and AuNPs for signal quantification. Figure 1 illustrates the design of a mAb-based blocking biosensor immunoassay for the detection of SARS-CoV-2 N protein specific antibodies in serum samples. Initially, PC will be prepared by coating viral N proteins on the surface and extra binding sites will be blocked by bovine serum albumin (BSA; Fig. 1B). Serum samples will be first added to the PDMS well and incubate for 1 hour. Subsequently, the reagent comprised of mAb-AuNPs will be introduced into the PDMS well (Fig. 1B), followed by PRAM imaging. In this simple 2-step assay, virus specific serum antibody (yellow) blocks the binding of mAb to viral protein (light blue) on the PC surface, while negative serum (no virus specific antibody) will allow the mAb to bind to the viral protein to form a AuNPs-mAb-antigen sandwich immunocomplex particle being captured onto the PC surface (Fig. 1C). The captured AuNPs locally quench the PC reflection efficiency. Peak intensity value image will be obtained by PRAM, which allows digital counting of particles for the quantitative analysis within 15 minutes. The reduction in AuNPs-labeled anti-N mAb binding to the antigen is directly proportional to the quantity of anti-N antibodies in the serum sample, providing an accurate quantitative measurement of serum antibody levels.

### Assessment of analytical sensitivity and limit of detection

Analytical sensitivity of blocking biosensor assay was determined by using positive and negative control serum standards from experimental cats. In this evaluation, we tested two-fold serial dilutions of the serum standards in duplicates. As presented in Figure 2, the dilution of 1:8 was the highest dilution that gives statistically significant difference between the positive and negative control standards, both at the AuNPs counts (Fig. 2A) and percentage inhibition (PI) ratios (Fig. 2B). To further determine the limit of detection (LOD), negative control standard cat serum was spiked with known concentrations of SARS-CoV-2 anti-N mAb and serially diluted in 2-fold. As shown in Figure 3A, there is an increase in the number of gold nanoparticles (AuNPs) bound to the surface through the antibody-antigen immunocomplex with the decreasing concentrations of SARS-CoV-2 anti-N mAb, which is consistent with the calculated PI values, with the limit of detection for anti-N mAb in cat serum determined to be 12.5 ng.

**Figure 2.**
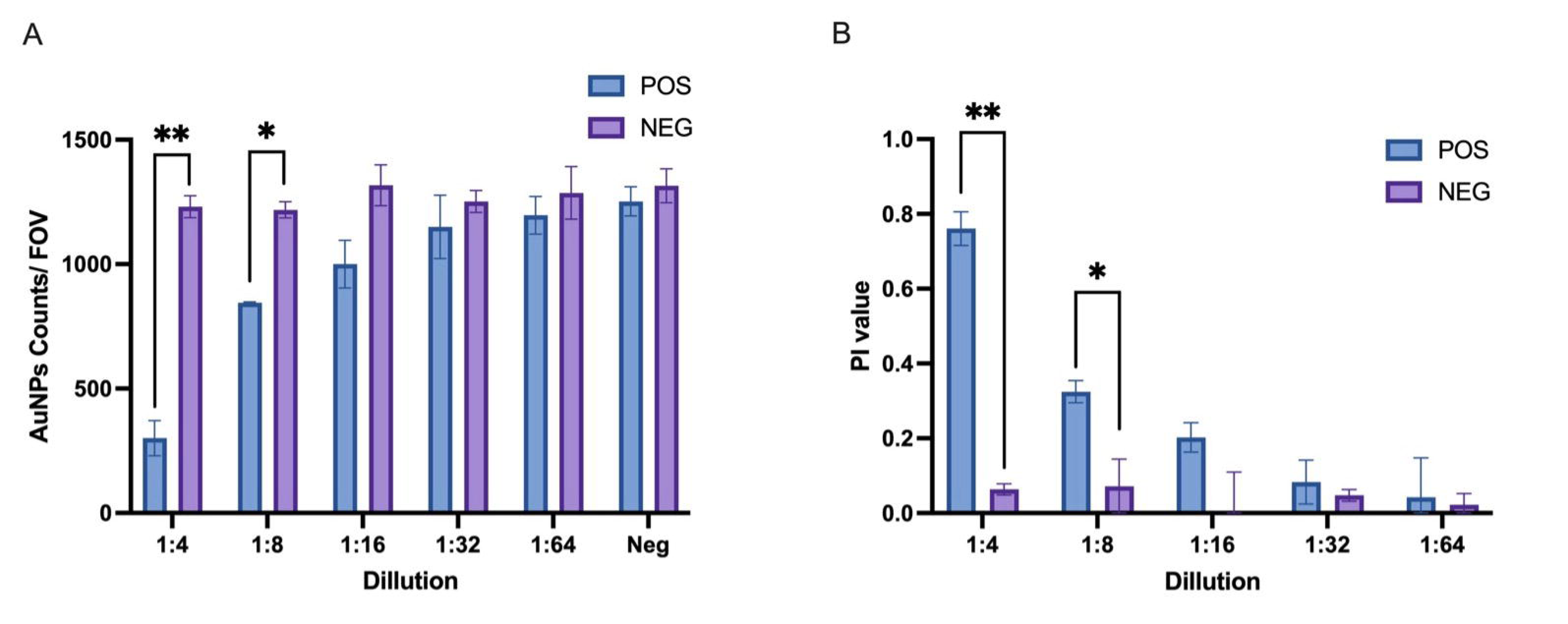
Analytical sensitivity of blocking biosensor assay in standard cat serum. Two-fold serial dilutions of the high-positive and negative control serum were tested in Blocking Biosensor assay. Each data point is shown as the mean value of two repeats, and the error bar represents the standard deviation. Asterisks indicate dilutions at which the AuNPs Counts/ FOV(A) and PI (B) of the positive control serum were statistically different from those of negative control serum. ( **p<0.01, *p<0.05).

**Figure 3.**
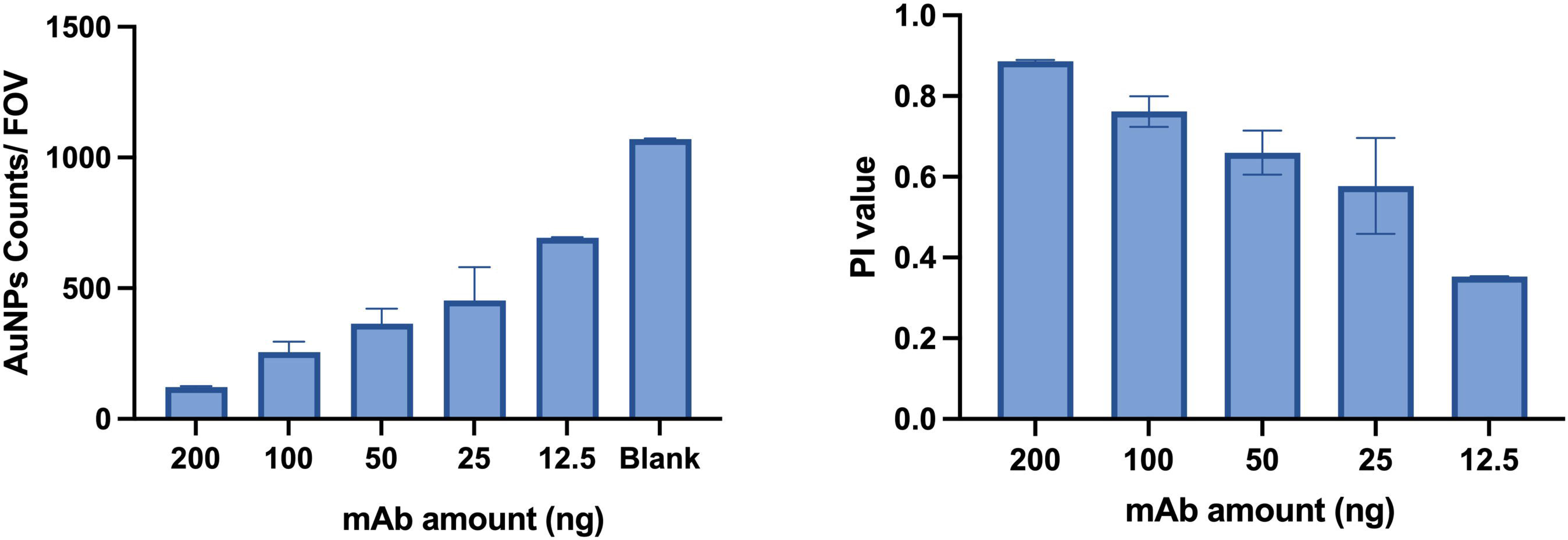
Limit of Detection (LOD) of blocking biosensor assay. (**A**) Particle quantification as a function of SARS-CoV-2 anti-N mAb amount in negative cats serum at 15 min. The error bars represent the standard deviation of two independent tests. (**B**) Corresponding PI values for the data presented in panel A.

### Determination of diagnostic sensitivity and specificity

To evaluate the precision of the blocking biosensor assay for antibody detection, we tested a cohort of serum samples with known antibody statuses, which includes 86 known positive and 76 known negative samples from experimental cats. These samples were confirmed antibody statuses in our previous studies^29^. As shown in Figure 4A, receiver operating characteristic (ROC) analysis revealed that at PI value cut-off at 0.5877, the assay achieved a diagnostic sensitivity of 97.80% and a diagnostic specificity of 98.67%. The area under the ROC curve (AUC) serves as an indicator of overall test accuracy, where an AUC of 1 represents perfect accuracy and an AUC above 0.9 is considered highly accurate. The AUC for this blocking biosensor assay is at 0.9983, with a statistical significance of P < 0.0001, demonstrating the high accuracy of the assay.

**Figure 4.**
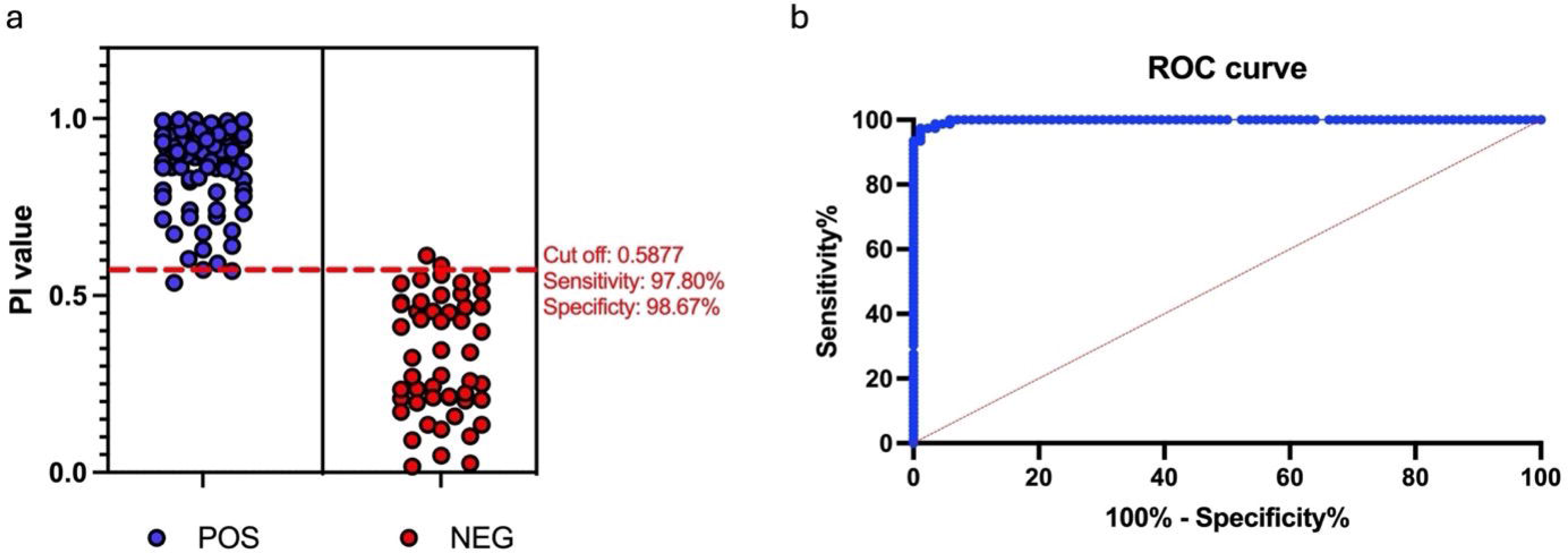
Determination of diagnostic sensitivity and specificity. Determination of diagnostic sensitivity and specificity was conducted through Receiver Operating Characteristic (ROC) analysis using 162 serum samples with known SARS-CoV-2 infection statuses (86 negatives and 76 positives). (A) An interactive plot displayed the cutoff value along with the optimal diagnostic sensitivity and specificity. (B) The ROC curve illustrated the accuracy, represented as the area under the curve (AUC). ROC analysis was carried out using GraphPad Prism version 10.0.0 for Windows, GraphPad Software, Boston, Massachusetts USA, www.graphpad.com.

### Blocking biosensor assay repeatability assessment

Repeatability in an assay refers to its consistency in producing the result when the same sample is prepared and tested multiple times. The repeatability of the blocking biosensor assay was evaluated using a single batch of positive control serum standard. To quantify the repeatability, the coefficient of variation percentage (%CV) was calculated. The results showed that the %CV within a single PC was 2.04% (mean value of 0.7073 ± standard deviation of 0.0144), the %CV between different PC in a single run was 2.73% (mean value of 0.7303 ± standard deviation of 0.02), and the %CV between different runs was 4.87% (mean value of 0.7150 ± standard deviation of 0.0348). These %CV values, bellowing 5%, indicate that the blocking biosensor assay is highly repeatable.

### Application of blocking biosensor assay in multiple animal species

To evaluate the cross-species assay performance of our blocking biosensor assay, we applied this assay in ferrets and deer samples with known antibody status. As shown in Figure 6, at the established cutoff PI value of 0.5877, the assay is able to distinguish between experimentally infected and negative control groups of animals. At 14 days post infection (DPI), all of the infected ferrets consistently showed PI values above the cutoff PI value (positive antibody status), while all 3 negative control ferrets remained below cutoff value (negative antibody status) (Fig. 5A). Similarly, at 21 DPI, all 5 infected deer were detected as positive for anti-N antibody, while all 3 negative control deer were detected as negative for anti-N antibody (Fig. 5B).

**Figure 5.**
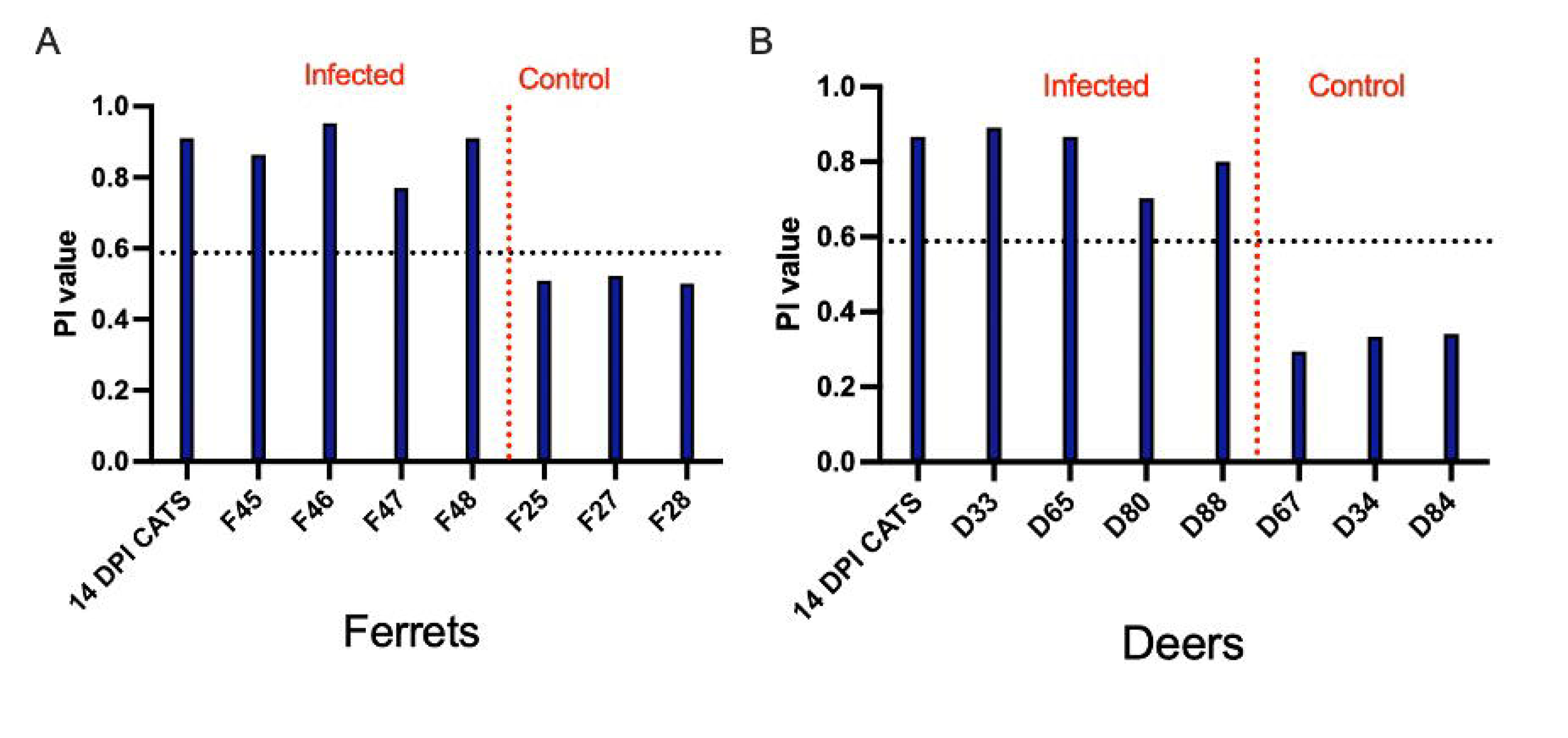
Detection of SARS-CoV-2 specific antibody responses in Ferrets and Deer. (A) Testing of ferret serum samples at 14 DPI; (**B**) Testing of deer serum samples at 21 DPI. Dotted line represents cutoff PI value of 0.5877. Both animals are infected with SARS-CoV-2 variant D416G.

**Figure 6.**
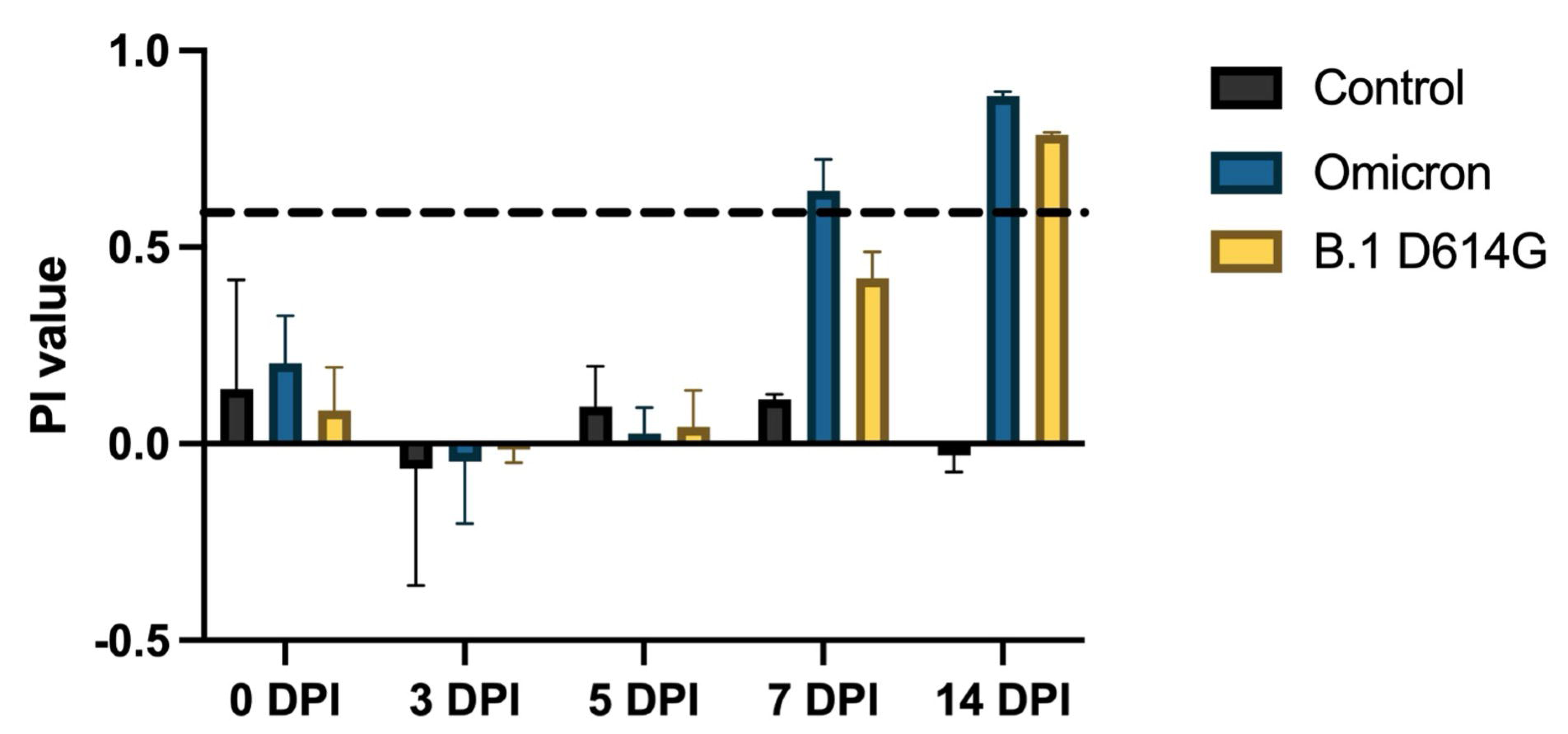
Dynamics of antibody response in domestic cats infected by SARS-CoV-2. Serum samples were collected from two cats infected with different SARS-CoV-2 variants at 0, 3, 7, 10, and 14 days post-inoculation, while samples from one negative control cat were used for comparison. The dashed line represents the cutoff PI value of the block biosensor assay. The x-axis indicates the time points of sample collection, while the y-axis displays the calculated percent inhibition (PI) value.

### Determine timeline of seroconversion in cats infected by SARS-CoV-2

We further applied our blocking biosensor assay to monitor the antibody response in experimental cats infected with the Omicron or B.1 D614G variants over the time course of infection. The result showed that SARS-CoV-2 specific antibody can be detected in animals infected by both variants at 14 DPI, while antibodies against Omicron variant can be detected as early as 7 DPI (Figure 6).

## DISCUSSION

In this study, we developed a novel mAb-based blocking biosensor assay. This is the first time integrated the blocking ELISA format into a biosensor system.

Compared to traditional bELISA, this biosensor assay has higher analytical and diagnostic sensitivity (98.3%). This result is expected because the biosensor system counts each antibody-antigen complex that is labelled with an AuNP tag, which is directly related to the number of antibody molecules from the test sample that compete for biosensor binding sites with AuNP-labeled anti-N mAb molecules that comprise the reagent included with the test sample.

In addition to the higher diagnostic sensitivity, the assay can reach diagnostic specificity of 96.5%. As indicated above, this biosensor assay has integrated the advantage of mAb-bELISA. In our previous study^29^, the mAb 127-3 was determined to specifically recognize a conserved epitope on SARS-CoV-2 N protein, but not cross-react with the N protein of other human/animal coronaviruses. On the other hand, since the biosensor reads the signal from mAb 127-3 conjugated AuNPs, no animal species specific secondary antibody is needed. This unique property makes our assay applicable to all animal species and humans. In this study, we demonstrated that this blocking biosensor assay is capable of detecting host antibody response against SARS-CoV-2 infection in multiple species, including cats, ferrets, and deer. Besides the diagnostic sensitivity and specificity assessment, repeatability of the assay was evaluated, resulting in sufficient lower coefficient of variation (less than 5%) across different trials, demonstrating the consistency and reliability of this test.

Compared to SARS-CoV-2 S protein, another commonly used antigen for antibody tests, the N protein region is more conserved among different SARS-CoV-2 variants, which makes this N-antigen base biosensor assay more suitable for detecting the exposure from different viral variants. Using this blocking biosensor assay, we were able to study the dynamics of antibody response in experimental cats infected by different SARS-CoV-2 variants. Specifically, we determined that the assay is able to detect anti-N antibody response in cats as early as 7 DPI for Omicron variant and 14 DPI for D614G variant. This has significant implications for early detection and monitoring of the coronavirus infection, particularly in diversified mammalian hosts.

In conclusion, this blocking biosensor assay provides a powerful diagnostic tool with increased sensitivity, specificity, and versatility necessary for managing the challenges posed by SARS-CoV-2 across various animal species. It not only enhances our diagnostic capability against current viral strains but also equips us to detect emerging variants of the virus, which is important for maintaining public health security and controlling the pandemic impact on both human and animal health.

## ACKNOWLEDGEMENT

This project was supported by NIH (grant #R01AI166791).

